# Microglial GSDMD-Mediated Pyroptosis Drives Neuroinflammation in Parkinson’s Disease

**DOI:** 10.1101/2025.06.30.662314

**Authors:** Mengmeng Wang, Yuedong Wang, Tianyi Wang, Yi Han, Tao Huang, Jicong Du, You Yin

## Abstract

**Rationale:** Parkinson’s disease (PD), a globally prevalent neurodegenerative disorder, is characterized by substantia nigra dopaminergic neuron degeneration and striatal dopamine depletion. While microglial pyroptosis is implicated in neuroinflammation and neural injury via inflammatory cytokine release, the role of the CASPASE-1/GSDMD pathway in PD pathogenesis remains incompletely defined.

**Methods:** 1-methyl-4-phenyl-1,2,3,6-tetrahydropyridine (MPTP) was used to construct PD model i*n vivo*, GSDMD-knockout mice was employed to assess pyroptotic mechanisms. MPP⁺-stimulated BV2 microglia were treated with a CASPASE-1 inhibitor *in vitro*. Microglia-specific GSDMD conditional knockout mice were generated to evaluate cell-type contributions to neuroinflammation and motor deficits.

**Results:** GSDMD deficiency attenuated MPTP-induced neuroinflammation, dopaminergic neuron loss, and motor dysfunction in vivo. MPP⁺ exposure triggered NLRP3 inflammasome activation and pyroptosis in BV2 microglia, which was suppressed by CASPASE-1 inhibition. Critically, microglia-specific GSDMD ablation mitigated nigrostriatal degeneration and dyskinesia in PD mice, confirming the centrality of microglial pyroptosis.

**Conclusion:** Our findings demonstrate that microglia drive neuroinflammation in PD via CASPASE-1/GSDMD-mediated pyroptosis, directly linking this pathway to dopaminergic neurodegeneration and motor impairment. Targeting GSDMD-dependent pyroptosis represents a promising therapeutic strategy.

## Introduction

Parkinson’s disease (PD) is the second most intractable neurodegenerative disease in the world. which brings heavy physical and economic burden to the aging population(Bloem, Okun et al., 2021). PD is more common in elderly patients, with a prevalence rate of 1%-2% among people over 65 years old(Bloem et al., 2021). PD is characterized by degeneration and death of dopaminergic neurons in the substantia nigra (SN), resulting in reduced dopamine levels in the striatum, which manifest as motor dysfunction, mainly contained slow movement, tremors, muscle stiffness, and other physical and involuntary movement disorders(Armstrong & Okun, 2020).

Conclusions from clinical cases and animal models have proved that the damage of dopaminergic neurons caused by neuroinflammation can occur at the early stage of PD. Elevated levels of pro-inflammatory cytokine were found in the SN of PD patients and animal model. Microglia induced neuroinflammation is the core event of in PD(Bartels, De Schepper et al., 2020). Activation of microglia may give rise to neuronal damage through the release of inflammatory cytokines and neurotoxic metabolites. Inhibition of neuronal damage mediated by microglia activation may provide a potential cure for alleviating PD progression. Activated microglia and increase of inflammasomes of NOD-like receptor thermal protein domain associated protein 3(NLRP3) were observed in the SN(Gordon, Albornoz et al., 2018). NLRP3 inflammasomes is a macromolecular multi protein complex consisting of NLRP3, effector protein CASPASE-1 and adaptor protein ASC(Han, Sun et al., 2019). NLRP3 inflammasomes-mediated neuroinflammation acts as a vital role in the process of dopaminergic neurons degeneration in PD(Bartels et al., 2020). In previous studies we found that neuroinflammation was highly correlated with the NLRP3 inflammasomes activated in microglia(Cheng, Liao et al., 2020, Haque, Akther et al., 2020, Wang, Yuan et al., 2019). The activation NLRP3 inflammasomes was the important reason for the degeneration of dopaminergic neurons *in vitro* and *in vivo* (Haque et al., 2020).

Pyroptosis, a new type of programmed cell death, is caused by the stress stimulation such as infection and injury. The Gasdermin protein family was identified as the key executive protein in the process of Pyroptosis. As an executive molecule, GSDMD protein consists of C-terminal domain, N-terminal domain and intermediate connection. C-terminal domain is water-soluble, which inhibits the cytotoxic effect of N-terminal. N-terminal domain is fat-soluble, which acts as the key part in executing Pyroptosis(Broz, Pelegrín et al., 2020). GSDMD is cleaved by caspases to form 22 kDa C-terminal domain and 31 kDa N-terminal domain. The N-terminal domain can target at the cell membrane and insert into it, forming the 16 polymers on the cell membrane. After that, non-selective membrane pore of about 10-15 nm in inner diameter come into being, leading to the release of cell contents and cell swelling and resulting in pyroptosis (Kesavardhana, Malireddi et al., 2020, Tummers & Green, 2022). After triggered by different cytoplasmic sensor proteins recognizing PAMPs or DAMPs, the NLRP3 inflammasomes recruited pro-Caspase-1 monomers through the adaptor protein ASC (apoptosis-associated speck-like protein containing CARD) and activate the Caspase-1 by dimerization, making it a fully functional protease (p20/p10). Activated Caspase-1 cleaves GSDMD to release the N-terminal domain, finally causing pyroptosis. Caspase-1 also processes the proinflammatory cytokine pro-IL-1β to generate mature IL-1β, which is presumably released by cell lysis during Pyroptosis,

In our study, we revealed GSDMD deficiency significantly alleviated motor dysfunction symptoms and reduced the degeneration of dopaminergic neurons *in vivo*. Moreover, CASPASE-1 deficiency significantly inhibited the level of neuroinflammation caused by MPTP *in vivo*. Meanwhile, GSDMD cleavage and neuroinflammation were also inhibited in CASPASE-1 KO mice. In addition, we found inflammasome activation and BV2 cells pyroptosis after MPP+ stimuli, which could be inhibited by CASPASE-1 inhibitor VX-765. By using conditional knockout mice, we proved GSDMD deficiency in microglia lead to mitigate PD dyskinesia symptoms and reduce degeneration of dopamine neurons. These results highlighted that microglia promote neuroinflammation *via* CASPASE-1/GSDMD mediated Pyroptosis in PD.

## Materials and Methods

### Chemicals and reagents

1-Methyl-4-phenyl-1,2,3,6-tetrahydropyridine (MPTP), 1-methyl-4-phenylpyridinium iodide (MPP^+^ iodide) and Belnacasan (VX-765) were purchased from Selleckchem chemicals. LEGENDplex™ Mouse Inflammation Panel (13-plex) was purchased from Biolegend. TransDetect Apoptosis Detection Kit was purchased from TransGen Biotech (Beijing, China). Polyclonal Antibody CASPASE-1 P10-FITC conjugated was purchased from Signalway Antibody (USA). The PCR kit was purchased from TAKARA (Japan). The primes for Q-PCR were purchased from Shenggong Biotech (Shanghai, China). PBS, DMEM, and fetal bovine serum (FBS) were purchased from Gibco (New York, USA). The antibodies (GAPDH, Cleaved CASPASE-1) were purchased from Cell Signaling Technology (Massachusetts, USA). The GSDMD antibody was purchased from ABCAM (Cambridge, UK).

### Mice and treatment

Male C57BL/6 mice, 6-8 weeks, were purchased from China Academy of Science (Shanghai, China). GSDMD KO and CASPASE-1 KO, GSDMD^flox/flox^ and CX3CR1^Cre^GSDMD^flox/flox^ mice 6-8 weeks, were purchased from Cyagen (Jiangsu, China). All mice were kept in a constant temperature, 12h light-dark cycle laboratory animal room with adequate food and water supplies. Mice were intraperitoneally injected with MPTP (30.0 mg/Kg, every 24 h for 5 days) to create PD models.

### Mice behavioral test

Behavioral tests were conducted 24 hours after the completion of the final PBS or MPTP injection. The mice were sent to the behavioral laboratory 1-2 hours early to acclimate.

#### Open field experiment

The mice started in the corner of an open-field device (500 x 500 x 300 mm) and explored spontaneously for 10 minutes under an overhead video tracking system. The total activity distance and immobility time in the middle region of the mice were calculated by the computer tracer analysis system.

#### Pole climbing experiment

Take a cylindrical stick that has a rough wooden ball as top about 50 cm long, whose bottom was placed in a mouse cage. The mouse starts with its head up on the ball at the top of the pole, the time was started when mouse climbed down to the pole with its head down, the time was stopped when mice climbed along the pole to the bottom (with its front paws landing on the ground), and this period was recorded as the climbing time. The shorter the climbing time of mice, the stronger the motor ability of mice.

#### Rotarod experiment

Before the experiment, the mice were trained for 5 minutes on the adaptability of a 4 r/min rotating rod, and after 30 minutes with resting, mice were formally tested. Mice were placed on the rotating rod rotating continuously at the acceleration of 0-60 r/min, and the residence time of mice on the rotating rod was recorded. The longest detection time was 5 minutes.

### Cell culture and treatment

Microglia (BV2) cells (monocyte macrophage line) was obtained from American Type Culture Collection and cultured in DMEM with 10% FBS at 37°C in a 5% CO2 humidified chamber. BV2 cells were treated with MPP^+^ or MPP^+^ and VX-765(40 μM) for 6 h.

### Flow Cytometry

After stimulated with MPP^+^ or MPP^+^+VX-765(40 μM), BV2 cells were digested with EDTA free trypsin and fixed by fixation buffer. Then BV2 cells were stained with PI-PE antibody and Polyclonal Antibody CASPASE-1 P10-FITC antibody for 20 min to detect the rate of pyroptosis. After establishing the MPTP induced PD model, LEGEND plex™ Mouse Inflammation Panel (13-plex) was used to detect the level of inflammation cytokines including IL-1β, IL-18, IL-17A and IL-27 in the mice serum according to the instructions.

### Western Blot analysis

The protein was extracted from BV2 cells by using a mammalian protein extraction reagent according to the manufacturer’s protocol after stimulated with MPP^+^ or MPP^+^ +VX-765 (40 μM), then analyzed by WB to detect GAPDH, Cleaved CASPASE-1, GSDMD. The secondary antibody (1:1000) was purchased from Cell Signaling Technology.

### Histological examination

Mice were sacrificed 24 h after the final administration of PBS or MPTP. The SN of PD mice were collected from mice and then fixed in 4% paraformaldehyde. Nissl Staining and Immunofluorescence staining were performed according to the manufacturer’s instructions. The sections were stained with primary antibodies: anti-Glial Fibrillary Acidic Protein (GFAP), anti-NESTIN, anti-α-Synuclein (α-Syn), anti-Tyrosine Hydroxylase (TH), anti-Iba-1 and anti-GSDMD. Fluorescence was quantified by Image J (National Institutes of Health, Bethesda, MD, USA).

### Statistical analysis

Data were expressed as means ± standard deviation (SD). Two-tailed Student’s t-test was used to analyze the differences between two groups. One-way ANOVA was employed to analyze the differences among three groups. The data were analyzed using GraphPad Prism 8.0.1.244 and SPSS ver. 19 software (IBM Corp, Armonk, New York, USA). P < 0.05 was considered statistically significant.

## Results

### 1. GSDMD deletion mitigated MPTP-induced PD symptoms by reducing degeneration of dopamine neurons in the SN

PD model was established by using MPTP. The mice motor activity was significantly impaired after MPTP treatment. MPTP-induced dyskinesia symptoms were significantly alleviated in GSDMD KO mice compared with WT (Figure 1A/B). In addition, MPTP-induced hindlimb gripping behavior was significantly reduced in GSDMD KO mice compared to WT mice (Figure 1C/D/E). Consistent with these observations, MPTP induced severe trunk dystonia in WT mice, but less so in GSDMD KO mice. The results of mice behavioral tests showed that GSDMD deficiency alleviated motor dysfunction in PD mice, suggesting that GSDMD deficiency significantly enhanced the motor function in MPTP-induced PD mice models. The SN of PD mice were taken to evaluate whether GSDMD deficiency could alleviate MPTP-induced dopamine neurons degeneration. Nissl bodies are abundant in neurons with strong metabolic function and serve as marker of neuronal status. The amount of Nissl bodies was decreased significantly in WT mice after MPTP treatment, and GSDMD deficiency salvaged the loss of Nissl bodies (Figure 1F/G). Deficiency of Tyrosine hydroxylase (TH), a key enzyme in DA biosynthesis, is a sign of PD progression. As predicted, TH-positive dopaminergic neurons in WT mice were significantly reduced compared with GSDMD KO mice (Figure 1H/I). The abnormal assemble of α-Synuclein (α-Syn) in dopaminergic neurons is the crucial pathological feature of PD diagnosis and α-Syn was upregulated in MPTP-induced PD models(Armstrong & Okun, 2020, Charvin, Medori et al., 2018). As shown in (Figure 1J/K), the amount of α-Syn in WT mice was significantly higher than that in GSDMD KO mice. Analyzing the above experimental results, we found that the regulation of GSDMD might provide a potential therapeutic strategy for alleviating the loss of dopamine neurons in SN induced by MPTP.

**Figure 1.**
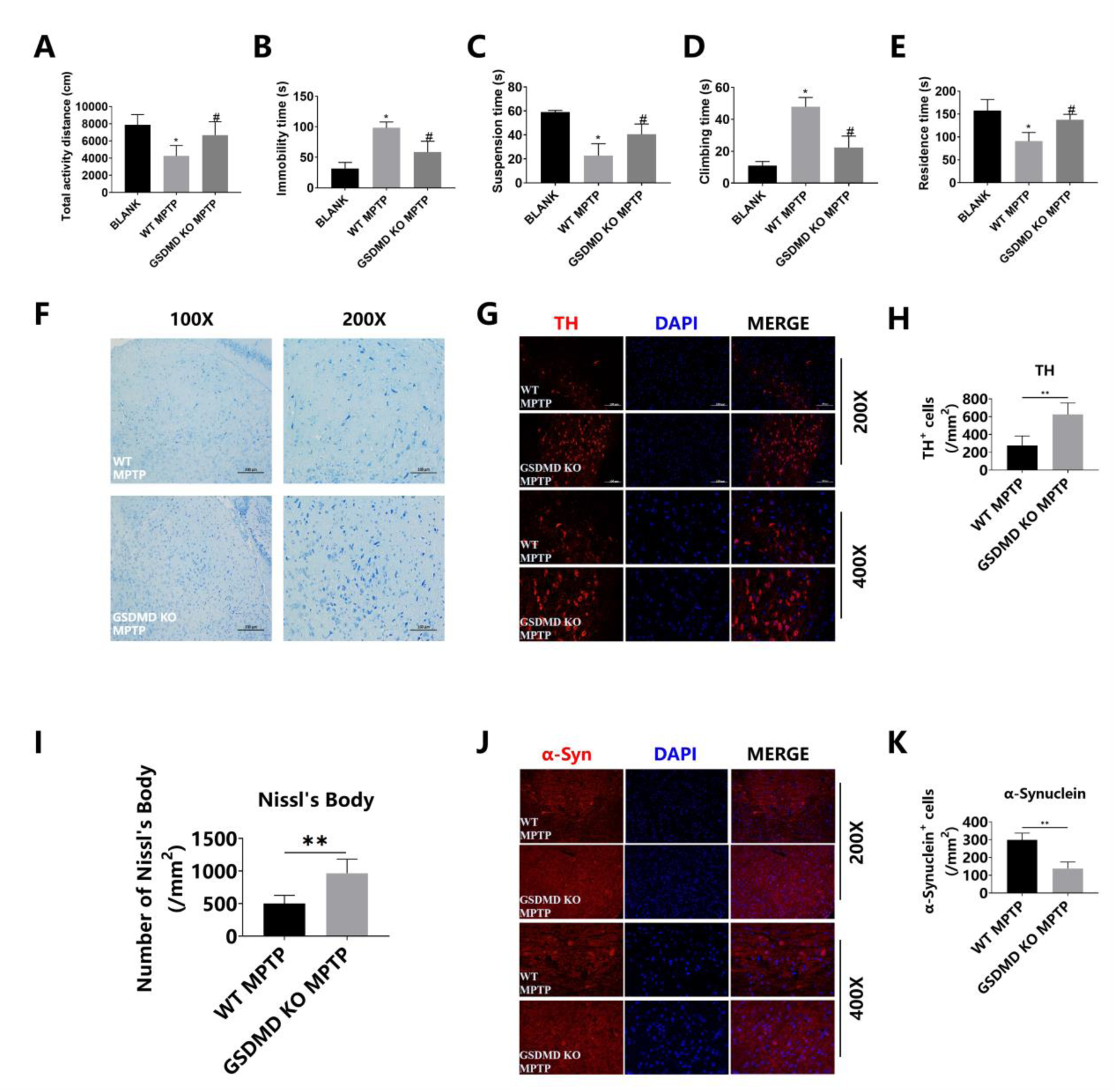
GSDMD deficiency attenuated MPTP-induced PD symptoms in the SN of MPTP-treated mice. WT and GSDMD KO mice were treated with PBS or MPTP for 5 days. **A.** Total activity distance of PD mice in open field experiments. **B.** The immobility time of PD mice in open field experiments. **C.** The suspension time of PD mice in suspension experiments. **D.** The climbing time of PD mice in pole climbing experiments. **E.** The residence time of PD mice in the Rotarod experiments. **F.** Nissl body staining in SN. **G.** The number of Nissl body(/mm^2^). **H.** TH immunofluorescence in SN. **I.** The number of TH^+^ cells (/mm^2^). **J.** α-Syn immunofluorescence in SN. **K.** The number of α-Syn^+^ cells(/mm^2^). Data are presented as mean ± SD. *, P < 0.05; **, P<0.01.

### 2. GSDMD deficiency attenuated neuronal damage and neuroinflammation

To observe whether GSDMD deficiency could alleviate MPTP-induced neuronal damage, the SN of mice were taken for immunofluorescence after MPTP treatment. Compared with GSDMD KO mice, WT mice showed a significant reduction in GFAP-positive neurons in SN (Figure 2A/B). Meanwhile, a significant reduction in NESTIN levels was observed in WT mice, which was not observed in GSDMD KO mice (Figure 2C/D). Dopaminergic neuron damage is often caused by early neuroinflammation in PD (Hammond, Marsh et al., 2019, Ising, Venegas et al., 2019). The level of neuroinflammation in PD mice was detected by using Flow Cytometry. The results showed that the levels of IL-1β, IL-18, IL-17A, IL-27 were increased after MPTP administration, while GSDMD deficiency significantly inhibited the production of these proinflammatory cytokines (Figure 2E). These results confirmed that GSDMD deficiency significantly alleviated neuroinflammation in PD. Next, QT-PCR was used to evaluated the level of neuroinflammation in the SN of PD mice. The results showed that the expression level of proinflammatory cytokines and chemokines including IL-1β, TNF-α, CCL5, CXCL9, CXCL10, and CXCL11 in GSDMD KO mice was obviously inferior to WT mice (Figure 2F). Taken together, these results indicated that GSDMD deficiency attenuated neuronal damage and neuroinflammation.

**Figure 2.**
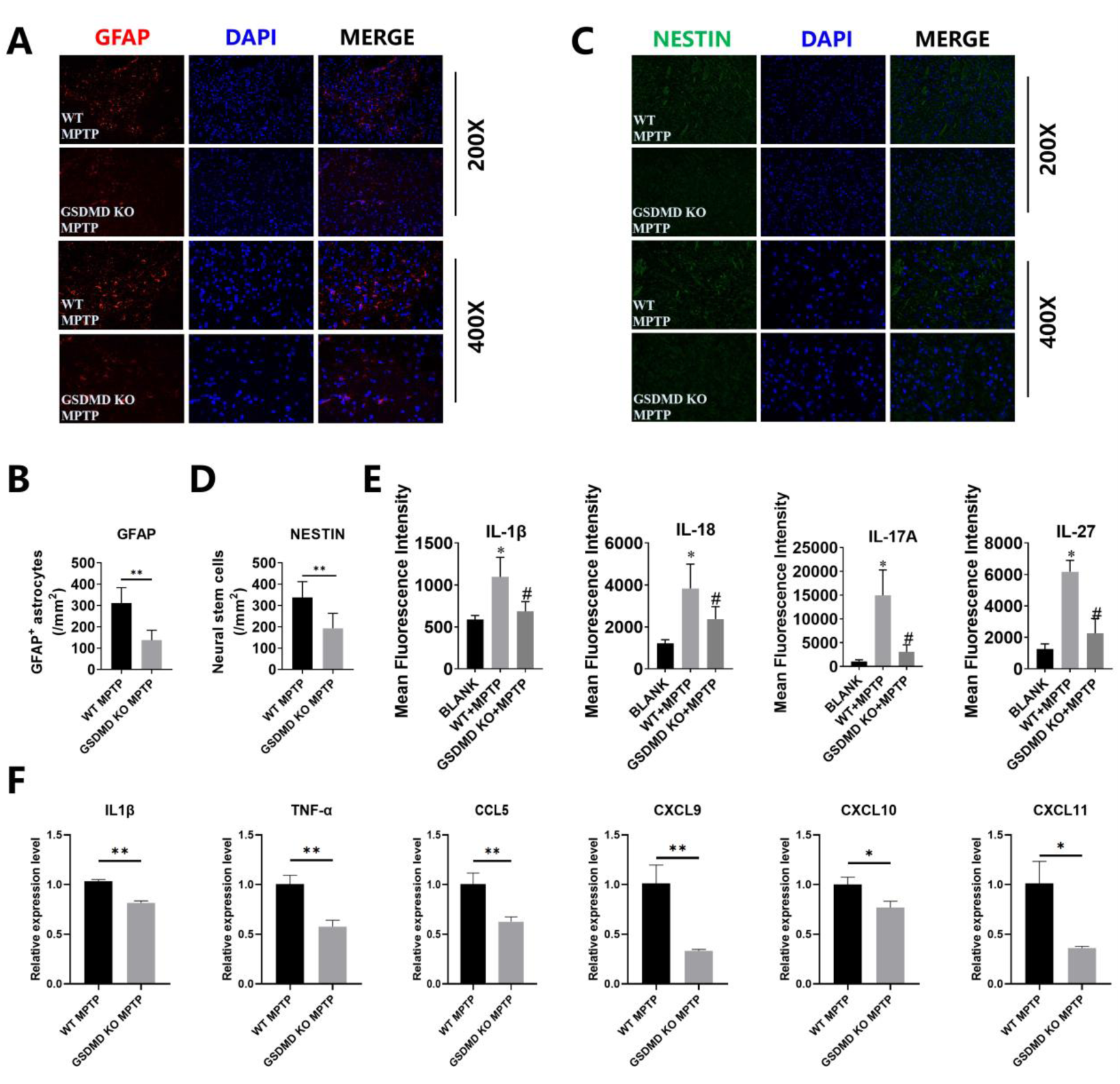
GSDMD deficiency attenuated microglia activation, neuronal damage and neuroinflammation levels. WT and GSDMD KO mice were treated with PBS or MPTP for 5 days. **A.** GFAP immunofluorescence in SN. **B.** Quantitative analysis and the statistics on the number of GFAP^+^ cells(/mm^2^). **C.** NESTIN immunofluorescence in SN. **D.** Quantitative analysis and the statistics on the number of NESTIN^+^ cells(/mm^2^). **E.** The levels of IL-1β, IL-18, IL-17A, IL-27 in mice serum. **F.** QT-PCR detection of RNA extracted from the SN of mice. Data are presented as mean ± SD. *, P < 0.05; **, P<0.01.

### 3. CASPASE-1 deficiency attenuated microglial pyroptosis in MPTP induced PD

BV2 cells were used to examine whether CASPASE-1 activation is responsible for microglial pyroptosis in PD. BV2 cells were treated with MPP^+^ at different concentration gradients (0.25 mM, 0.5 mM and 1.0 mM) for 6 h. Then the levels of cleaved CASPASE-1 and N-GSDMD were detected. The WB results revealed the levels of cleaved CASPASE-1 and N-GSDMD were increased with gradient concentration (Figure 3A). Next, the pyroptosis rate of BV2 cells was evaluated by using Flow Cytometry. The pyroptosis rate of BV2 cells was increased after MPP^+^ stimulation (Figure 3B). Then we administered BV2 cells with CASPASE-1inhibitor and found the levels of cleaved CASPASE-1 and N-GSDMD were significantly decreased in VX-765 group (Figure 3C). The pyroptosis rate of BV2 cells was also decreased after VX-765 treatment (Figure 3D). Moreover, CASPASE-1 KO mice were used to discuss the function of CASPASE-1 in MPTP induced microglial pyroptosis in PD mice model. Notably, MPTP treatment promoted microglia pyroptosis, as marked by Iba-1 and GSDMD, in the SN of WT mice, but not in CASPASE-1 KO mice (Figure 3E/F). Taken together, these results indicated that CASPASE-1 deficiency attenuated MPTP-induced microglial pyroptosis and GSDMD mediated microglial pyroptosis was CASPASE-1 dependent.

**Figure 3.**
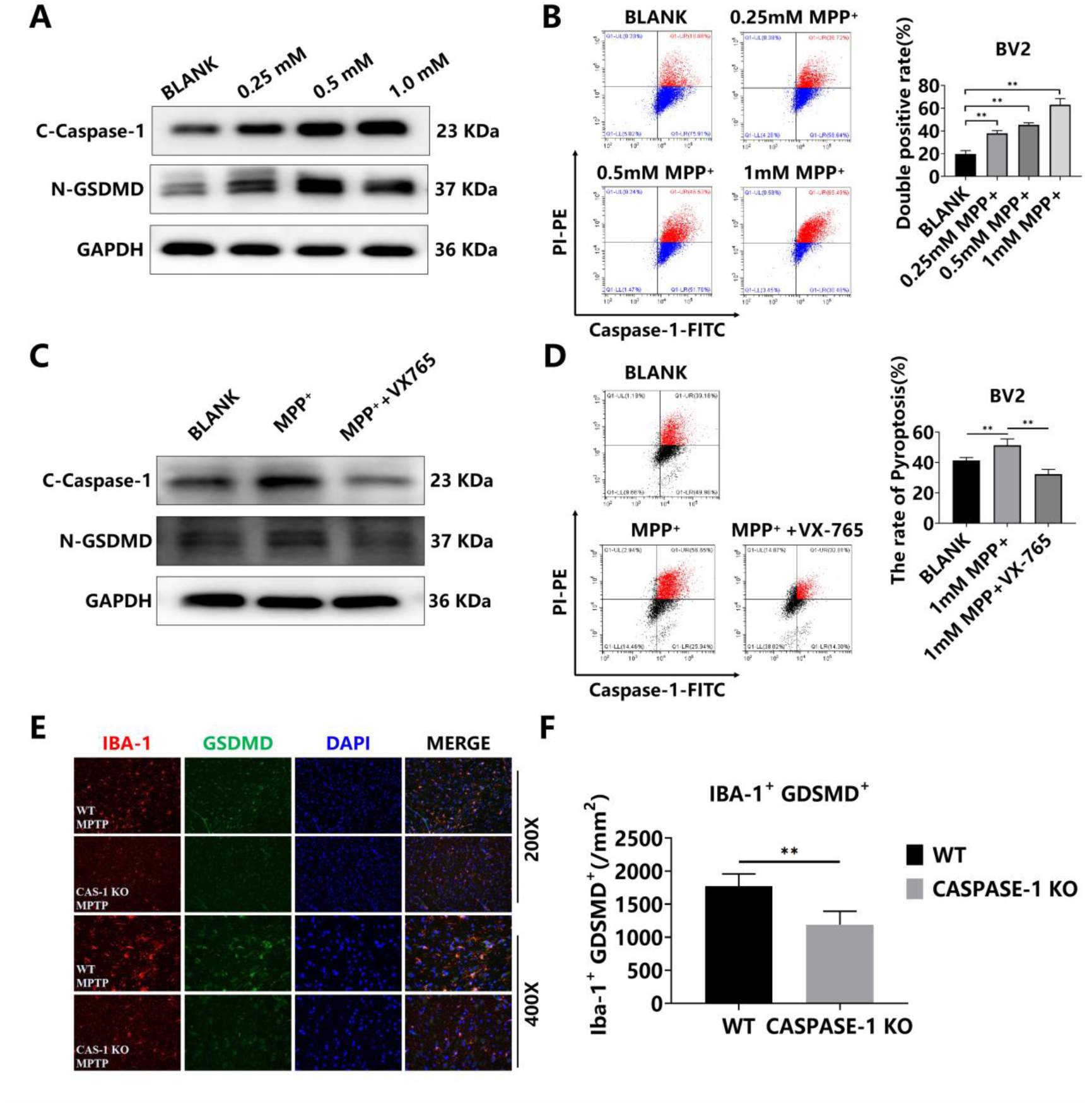
CASPASE-1 deficiency attenuated MPTP-induced microglial pyroptosis. **A.** WB analysis of BV2 cells: BLANK, 0.25 mM, 0.5 mM, 1.0 mM MPP^+^. **B.** Pyroptosis was detected by Flow Cytometry: BLANK, 0.25mM, 0.5mM, 0.5mM, 1mM MPP^+^. **C.** WB analysis of BV2 cells: BLANK, 1.0 mM MPP^+^, 1.0 mM MPP^+^+VX-765. **D.** Pyroptosis was detected by Flow Cytometry: BLANK, 1.0 mM MPP^+^, 1.0 mM MPP^+^+VX-765. **E.** Iba-1 and GSDMD staining in the SN of PD mice model. **F.** Quantitative analysis and the statistics on the number of Iba-1^+^ GDSMD^+^ cells (/mm^2^). Data are presented as mean ± SD. *, P < 0.05; **, P<0.01.

### 4. CASPASE-1 deficiency attenuated MPTP-induced PD symptoms by reducing degeneration of dopamine neurons in the SN

WT mice and CASPASE-1 KO mice were used to establish MPTP induced mice PD model. The results indicated that the motor function of CASPASE-1 KO mice was significantly better than that of WT mice. Through Open field experiments, Pole climbing experiments, and Rotarod experiments, we found the motor ability of CASPASE-1 KO mice was significantly stronger than that of WT mice (Figure 4A/B/C/D/E), indicating CASPASE-1 deficiency indeed attenuated motor dysfunctions in MPTP induced PD mice model. Next, Nissl bodies staining was performed to observe the state of neurons in the SN. The results showed that CASPASE-1 deficiency salvaged the loss of Nissl bodies in PD mice (Figure 4F/G). And the number of TH positive cells in SN region of CASPASE-1 KO mice was increased (Figure 4H/I), suggesting that CASPASE-1 deficiency significantly inhibited the damage of dopaminergic neurons in SN. Then we observed whether CASPASE-1 deficiency could attenuate abnormal aggregation of α-Syn in SN. The immunofluorescence result showed the number of α-Syn in CASPASE-1 KO mice was significantly lower than that of WT mice (Figure 4J/K). Taken together, these results indicated that CASPASE-1 deficiency attenuated MPTP-induced PD symptoms by reducing degeneration of dopamine neurons in the SN.

**Figure 4.**
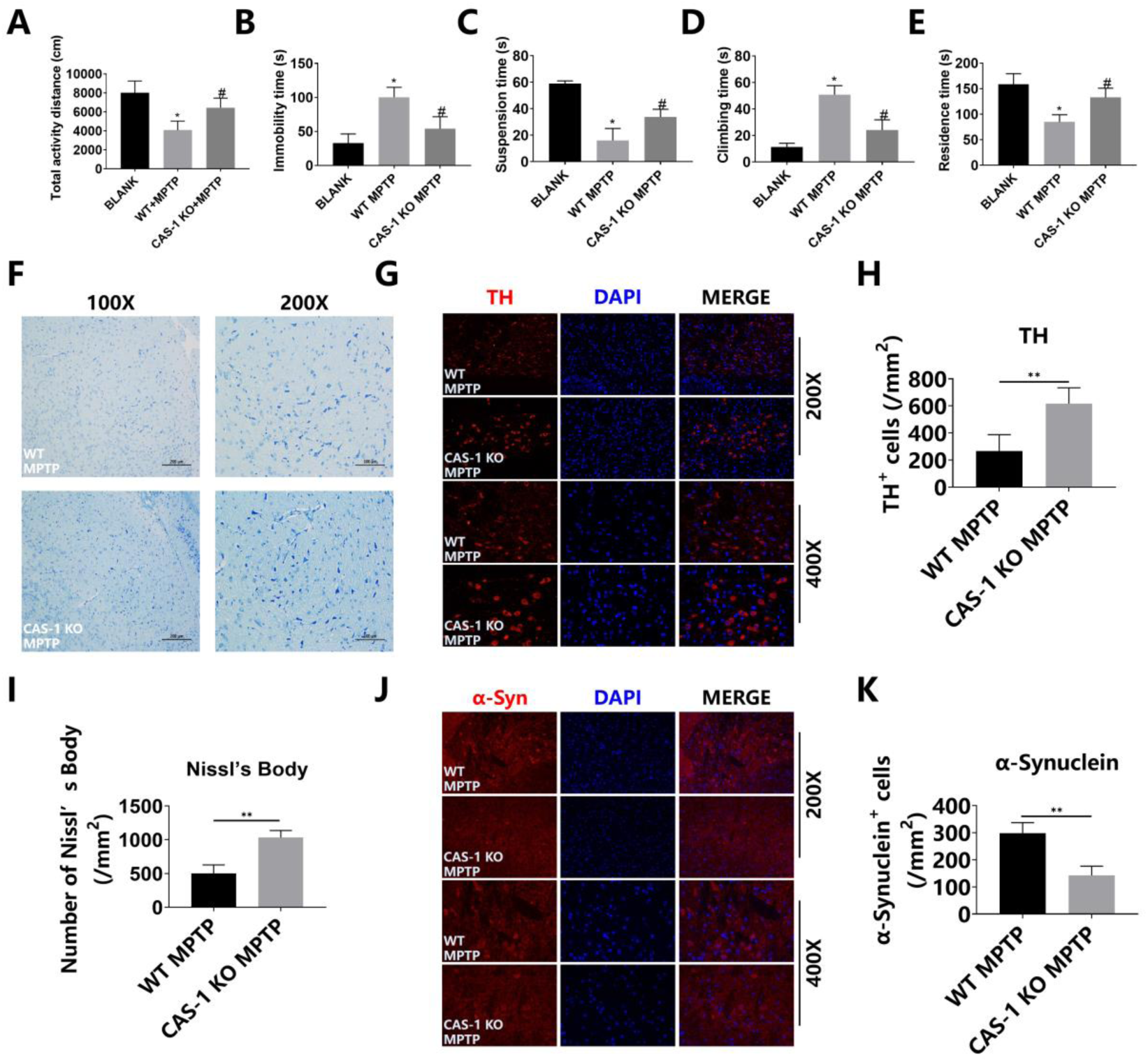
CASPASE-1 deficiency attenuated MPTP-induced PD symptoms in the SN of MPTP-treated mice. WT and CASPASE-1 KO mice were treated with PBS or MPTP for 5 days. **A.** Total activity distance of PD mice in open field experiments. **B.** The immobility time of PD mice in open field experiments. **C.** The suspension time of PD mice in suspension experiments. **D.** The climbing time of PD mice in pole climbing experiments. **E.** The residence time of PD mice in the Rotarod experiments. **F.** Nissl body staining in SN. **G.** Quantitative analysis and the statistics on the number of Nissl body(/mm^2^). **H.** TH immunofluorescence in SN. **I.** Quantitative analysis and the statistics on the number of and TH^+^ cells (/mm^2^). **J.** α-Syn immunofluorescence in SN. **K.** Quantitative analysis and the statistics on the number of α-Synuclein^+^ cells (/mm^2^). Data are presented as mean ± SD. *, P < 0.05; **, P<0.01.

### 5. CASPASE-1 deficiency attenuated neuronal damage and neuroinflammation

The SN of the mice brain was taken for immunofluorescence staining to observe whether CASPASE-1 deficiency could alleviate MPTP-induced neuronal damage by using WT and CASPASE-1 KO mice in PD model. The expression levels of GFAP (Figure 5A/B) and NESTIN (Figure 5C/D) in the SN of CASPASE-1 KO mice were significantly reduced compared to WT mice, which showed that pathological damage induced by MPTP in CASPASE-1 KO mice was alleviated. Moreover, the mice serum was extracted for detecting inflammation cytokines. We found inflammation cytokines in the serum of CASPASE-1 KO mice were down-regulated including IL-1β, IL-18, IL-17A, and IL-27, indicating that CASPASE-1 deficiency alleviated neuroinflammation in PD (Figure 5E). In addition, QT-PCR results showed the expression of IL-1β, TNF-α, CCL5, CXCL9, CXCL10 and CXCL11 were also obviously declined in CASPASE-1 KO mice, but not in WT mice (Figure 5F). Based on above results, it was confirmed that CASPASE-1 deficiency attenuated neuronal damage and neuroinflammation.

**Figure 5.**
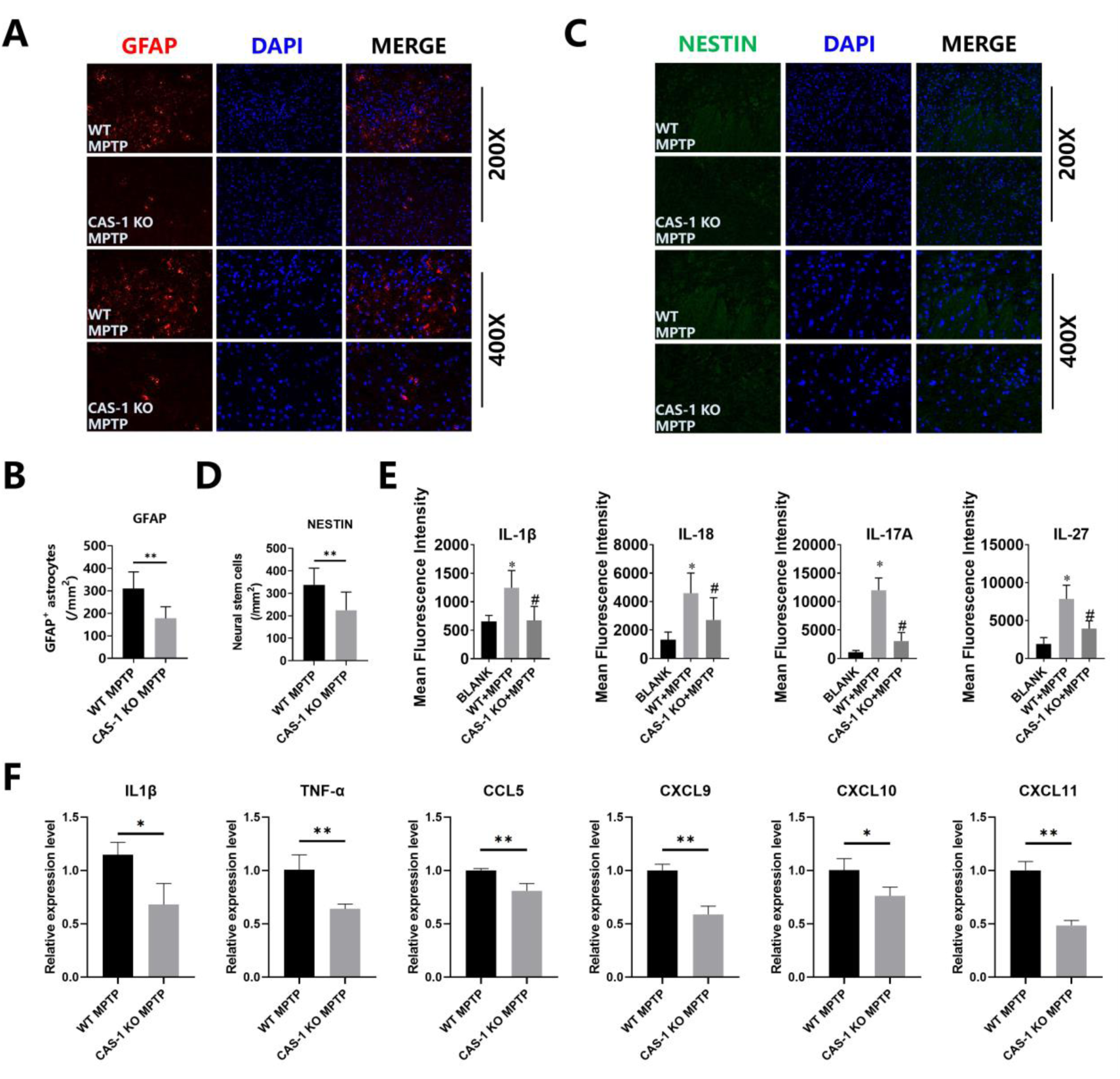
CASPASE-1 deficiency attenuated neuronal damage and neuroinflammation. WT and CASPASE-1 KO mice were treated with PBS or MPTP for 5 days. **A.** GFAP immunofluorescence in SN. **B.** Quantitative analysis and the statistics on the number of GFAP^+^ cells (/mm^2^). **C.** NESTIN immunofluorescence in SN. **D.** Quantitative analysis and the statistics of NESTIN^+^ cells number (/mm^2^). **E.** Serum cytokine levels in mice including IL-1β, IL-18, IL-17A, IL-27. **F.** QT-PCR detection of RNA extracted from the SN of PD mice. Data are presented as mean ± SD. *, P < 0.05; **, P<0.01.

### 6. GSDMD in microglia promoted PD dyskinesia symptoms and degeneration of dopamine neurons

To further evaluate the functions of GSDMD in microglia, we specifically knocked out GSDMD in microglia by expressing Cre recombinase under the control of the CX3CR1 promoter (CX3CR1^Cre^). To determine GSDMD functions in microglia, GSDMD^flox/flox^ and CX3CR1^Cre^GSDMD^flox/flox^ mice were injected with MPTP. We found the deletion of GSDMD in microglia alleviated motor dysfunction in PD mice (Figure 6A-F), indicating that loss of microglial GSDMD was sufficient to mitigate PD dyskinesia symptoms. To determine whether deficiency of GSDMD could reduce degeneration of dopamine neurons, we immunostained brain SN sections with Th and α-Syn antibodies. And the number of TH positive cells in SN region of CX3CR1^Cre^GSDMD^flox/flox^ mice was increased (Figure 8G/I), suggesting that GSDMD deficiency in microglia significantly inhibited the damage of dopaminergic neurons in SN. Then we observed whether GSDMD deficiency in microglia could attenuate abnormal aggregation of α-Syn in SN. The result of α-Syn immunofluorescence showed that the number of α-Syn in CX3CR1^Cre^GSDMD^flox/flox^ mice was significantly lower than that of WT mice (Figure 6H/J). Taken together, these results indicated that GSDMD in microglia promoted PD dyskinesia symptoms and degeneration of dopamine neurons.

**Figure 6.**
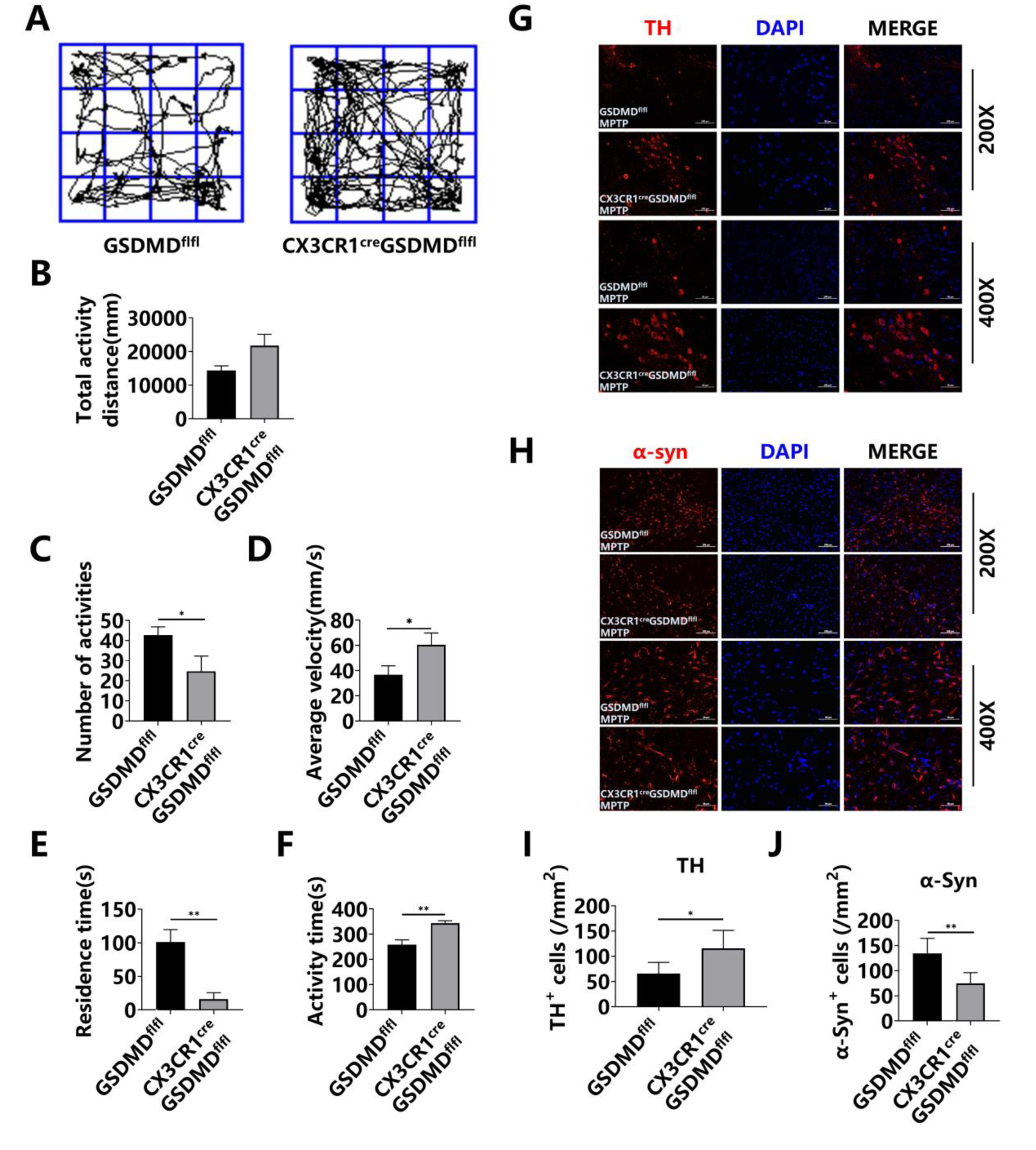
GSDMD in microglia promoted PD dyskinesia symptoms and degeneration of dopamine neurons. GSDMD^flox/flox^ and CX3CR1^Cre^GSDMD^flox/flox^ mice were treated with MPTP for 5 days. **A.** Open field experiment. **B.** Total activity distance. **C.** Number of activities. **D.** Average velocity. **E.** Residence time. **F.** Activity time. **G.** TH immunofluorescence in SN. **H.** α-Syn immunofluorescence in SN. **I.** The number of TH^+^ cells. **J.** The number of α-Syn ^+^ cells. Data are presented as mean ± SD. *, P < 0.05; **, P<0.01.

### 7. GSDMD deficiency in microglia inhibited MPTP induced neuroinflammation

To observe whether GSDMD deficiency in microglia could alleviate MPTP-induced neuronal damage, the SN of the mice brain was taken for immunofluorescence staining to observe neuronal damage. The expression level of GFAP (Figure 7A/B) in the SN of CX3CR1^Cre^GSDMD^flox/flox^ mice were significantly reduced compared to GSDMD^flox/flox^ mice, which showed that pathological damage induced by PD in CX3CR1^Cre^GSDMD^flox/flox^ mice was alleviated. Moreover, the mice serum was extracted for detecting inflammation cytokines. We found that inflammation cytokines in the serum of CX3CR1^Cre^GSDMD^flox/flox^ mice were down-regulated including IL-17A, IL-1β, IL-6, IL-1α, and MCP-1 (Figure 7C/D/E/F), indicating that microglial-specific ablation of GSDMD alleviated the level of neuroinflammation in PD. In PD, cytotoxic CD8^+^ T cell infiltration has been reported in post-mortem brain tissues. CCL5 and CXCL9 are associated with CD8^+^ T cells infiltration. QT-PCR results showed GSDMD deficiency in microglia inhibited the expression of CCL5, CXCL9 (Figure 7G/H). In addition, QT-PCR results showed the expression of CD8A, IFN-γ, GMZB, and TNF-α were obviously declined in CX3CR1^Cre^GSDMD^flox/flox^ mice (Figure 7I-L). In addition, MPTP treatment promoted cytotoxic T cells infiltration, as marked by CD8 and GMZB, in the SN of GSDMD^flox/flox^ mice, but not in CX3CR1^Cre^GSDMD^flox/flox^ mice (Figure 7M/N), which reminded us GSDMD deficiency in microglia inhibited CD8^+^ T cells infiltration and cytotoxicity. Based on above results, it was confirmed that GSDMD deficiency in microglia mitigate neuroinflammation.

**Figure 7.**
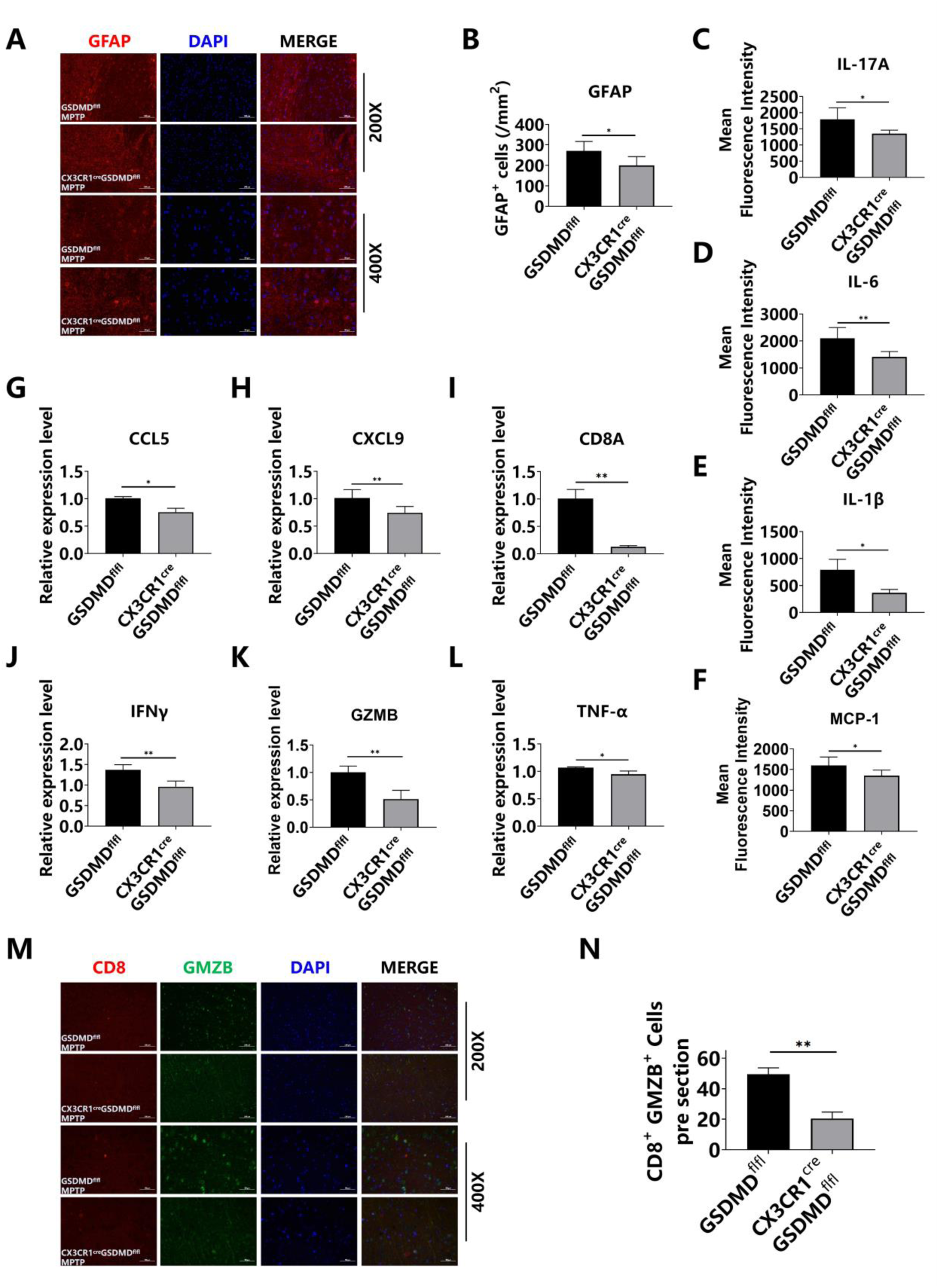
GSDMD deficiency in microglia inhibited MPTP induced neuroinflammation. GSDMD^flox/flox^ and CX3CR1^Cre^GSDMD^flox/flox^ mice were treated with MPTP for 5 days. **A.** GAFP immunofluorescence in SN. **B.** The number of GFAP ^+^ cells. **C-F.** Serum cytokine levels in mice including IL-17A, IL-1β, IL-6, IL-1α, and MCP-1. **G-L.** QT-PCR detection of RNA extracted from the dense part of the SN of PD mice. **M.** CD8 and GMZB staining in the SN of PD mice model. **N.** Quantitative analysis and the statistics on the number of CD8^+^ GMZB^+^ cells. Data are presented as mean ± SD. *, P < 0.05; **, P<0.01.

**Figure 8.**
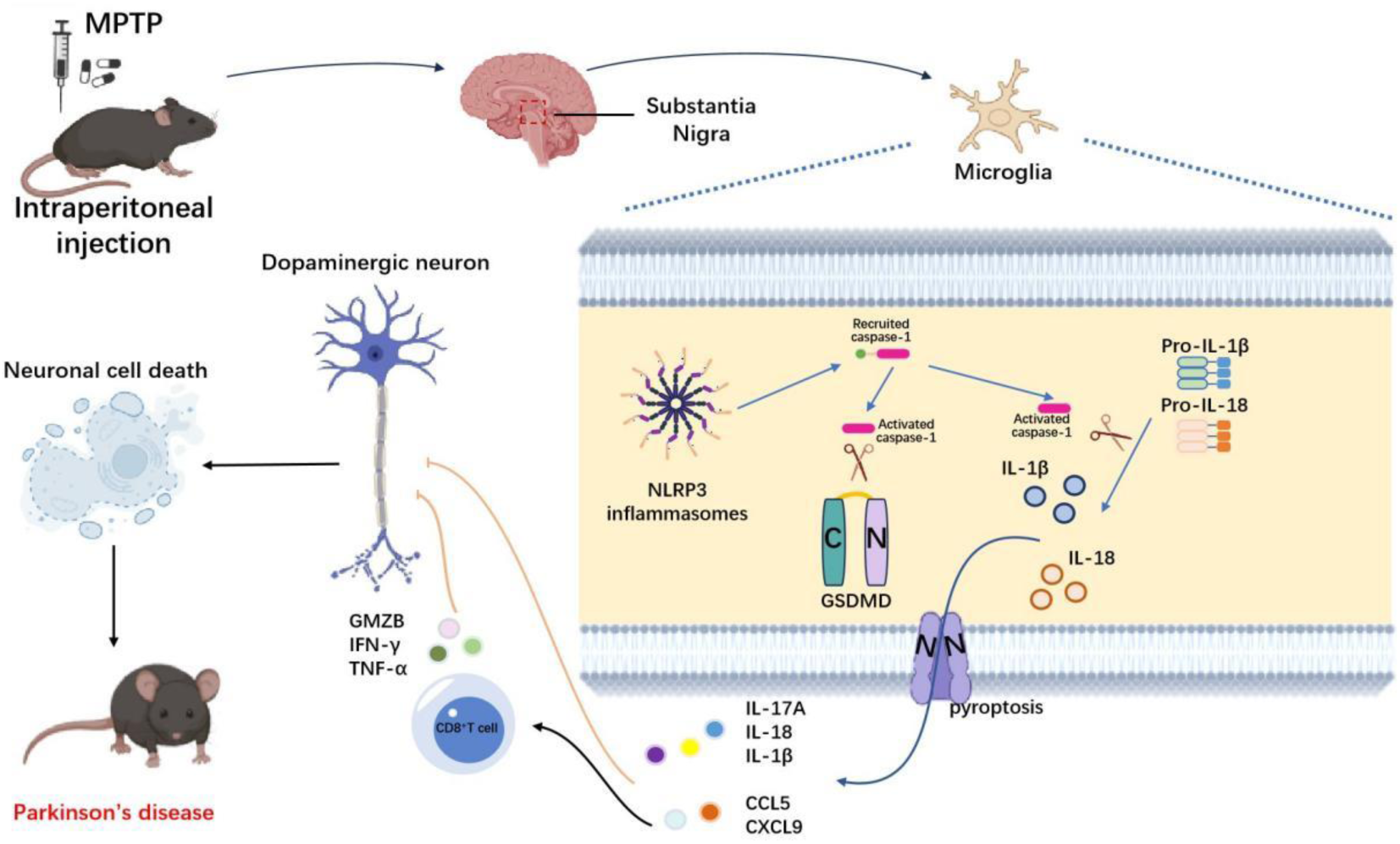
Microglia promote neuroinflammation via GSDMD mediated Pyroptosis in Parkinson’s disease.

## Discussion

PD is a common neurodegenerative disease that is significantly associated with age and is typically characterized by the loss of dopaminergic neurons(Tolosa, Garrido et al., 2021). PD is related to a variety of factors, such as age, lifestyle and environmental factors. The decrease of dopaminergic neurons in the dense part of SN and the abnormal assemble of α-Synuclein dopaminergic neurons are the main pathological feature of PD(Armstrong & Okun, 2020, Charvin et al., 2018). Therefore, current PD drugs are mainly aimed at supplementing exogenous dopamine and activation of dopamine receptors. But these drugs cannot prevent the progression of PD(Charvin et al., 2018, Parmar, Grealish et al., 2020, Vijiaratnam, Simuni et al., 2021). It is necessary to conduct researches on the mechanism of PD in order to restrain the progression of PD efficaciously.

Researches revealed that NLRP3 inflammasomes induced neuroinflammation plays roles in the regression of dopaminergic neurons. The activation of microglia is the core event of PD neuroinflammation. For example, Zheng Ruimao et al. found that celastrol, as a neuroprotective agent, could inhibit the activation of NLRP3 inflammasomes, and alleviate dopaminergic degeneration(Zhang, Zhao et al., 2021). Hu Gang et al. found small molecule kaempferol (Ka) could inhibit the activation of NLRP3 inflammasomes by reducing cleave CASPASE-1, and ultimately alleviate the LPS induced the loss of dopaminergic neurons(Han et al., 2019). These reports proved NLRP3 inflammasomes induced neuroinflammation in microglia may be a new target to prevent PD. In this work, our exploration emphasized the mechanism of PD progression that is a microglial functional pathway on the CASPASE-1-GSDMD-Pyrpptosis axis.

NLRP3 inflammasomes are characterized by NLRP3, ASC and CASPASE-1. The present of NLRP3 inflammasome has been demonstrated in PD and AD. NLRP3 activation can promote the dissociation of caspase-1 and accelerate the release of IL-1β and IL-18, thereby inducing pytoptosis. In 2015, GSDMD was affirmed as a crucial biomolecule in pyroptosis, which maked regulating pyroptosis as an effective way to regulate inflammatory diseases(Broz et al., 2020, Kayagaki, Stowe et al., 2015, Shi, Zhao et al., 2015). Recently, many studies showed that Gasdermin played important roles in tumors, inflammatory bowel disease and other diseases. Shao Feng et al. point out chemotherapeutic drugs can induce pyroptosis by cleaving GSDME through CASPASE-3, suggesting that regulating pyroptosis may be an efficient method to control the side-effect of chemotherapy(Wang, Gao et al., 2017), and they found that controllable activation of pyroptosis in situ of tumor can induce anti-tumor immunity (Wang, Wang et al., 2020). Liu Xing et al. reported that streptococcus pyogenes triggered pyroptosis of skin epithelium by cleaving GSDMA(Deng, Bai et al., 2022). Pyroptosis also play important role in neurological disease including PD. Christopher Powera et al. found that in multiple sclerosis, GSDMD-mediated pyroptosis has an positive effect(McKenzie, Mamik et al., 2018). Zheng Yuanyi et al. indicated Prussian Blue Nanozyme can reduce neurodegeneration by inhibiting pyroptosis(Ma, Hao et al., 2022). These studies strongly suggested that GSDMD played important roles in PD, but these studies only regard it as the most downstream molecule to execute pyroptosis, and there is still no direct study on the relationship between GSDMD and PD.

In this study, we found that GSDMD deficiency in microglia mitigated the PD symptoms. Firstly, we discussed the relationship between GSDMD and PD. Firstly, GSDMD KO mice and WT mice were used to established PD mice model. The results of mice behavioral test revealed that GSDMD deficiency attenuated motor dysfunctions in PD mice, which suggested that GSDMD played a critical role in MPTP-induced PD mice model. Next, Nissl body staining, α-Syn staining and TH staining were conducted to observe whether GSDMD deficiency could alleviate MPTP-induced PD symptoms. According to the immunofluorescence results, we discovered that modulating GSDMD could be a potential therapeutic method for alleviating MPTP induced PD symptoms. Notably, the reduction of dopaminergic neurons was significantly inhibited in the GSDMD KO group. Dopaminergic neuron damage is often caused by early neuroinflammation in PD. Research showed that the activation of microglia and the upregulation of pro-inflammatory cytokines such as TNF-α and IL-1β were the hallmarks of PD(Collins, Toulouse et al., 2012). And microglia activation was the core event of PD neuroinflammation(Hammond et al., 2019, Ising et al., 2019). Next, QT-PCR results showed that the levels of proinflammatory cytokines and chemokines including IL-1β, TNF-α, CCL5, CXCL9, CXCL10, and CXCL11 in GSDMD KO mice was significantly lower than that of WT mice. These results showed that GSDMD deficiency attenuated activation of microglia and the resulting neuronal damage and neuroinflammation levels.

In the progression of PD, the activation of microglia is the core event of PD neuroinflammation(Hammond et al., 2019, Ising et al., 2019). Next, BV2 cells were used to uncover the underlying molecular mechanisms underlying the contribution of microglial pyroptosis *in vitro*. BV2 cells were stimulated with different concentrations of MPP+. WB results prompted that the levels of N-GSDMD and activated-CASPASE-1 increased with increasing concentration of MPP+. But the level of N-GSDMD and pyroptosis rate were significantly decreased after VX-765 treatment. Moreover, Iba-1 and GSDMD immunofluorescence results showed that the level of cleaved GSDMD was significantly decreased in microglia of CASPASE-1 KO mice, indicating that CASPASE-1 deficiency attenuated MPTP-induced microglial pyroptosis. The results of mice behavioral test revealed that CASPASE-1 deficiency attenuated motor dysfunctions in PD mice. Similarly, CASPASE-1 deficiency also salvaged the loss of Nissl bodies in the SN. By α-Synuclein staining, it was found that the number of α-Syn in CASPASE-1 KO mice was significantly reduced. The expression of GFAP and NESTIN in the SN of CASPASE-1 KO mice were significantly reduced, but not in WT mice. In addition, QT-PCR results showed IL-1β, TNF-α, CCL5, CXCL9, CXCL10 and CXCL11 were significantly down-regulated in CASPASE-1 KO mice compared with WT mice. Taken together, these results revealed that CASPASE-1 deficiency attenuated MPTP-induced neuronal damage, microglia activation, and neuroinflammation.

At last, CX3CR1CreGSDMDflox/flox mice were used to figure out whether GSDMD deficiency in microglia could mitigate PD symptoms. The results showed that GSDMD deficiency in microglia attenuated PD symptoms and neuronal damage. Moreover, we found that inflammation cytokines in CX3CR1CreGSDMDflox/flox mice were down-regulated. In PD, cytotoxic CD8 T-cell infiltration has been reported in post-mortem brain tissues [Capelle CM., et al, Williams-Gray., et al]. CCL5 and CXCL9 are associated with CD8+ T cells infiltration [Dangaj D., et al, Hsu CL., et al]. QT-PCR results showed GSDMD deficiency in microglia inhibited the expression of CCL5, CXCL9. In addition, QT-PCR results showed the expression of CD8, IFN-γ, GMZB, and TNF-α were obviously declined in CX3CR1CreGSDMDflox/flox mice, which reminded us GSDMD deficiency in microglia inhibited CD8+ T cells infiltration and cytotoxicity. Based on above results, it was confirmed that GSDMD deficiency in microglia mitigate neuroinflammation.

In conclusion, GSDMD deficiency in microglia significantly inhibited the level of neuroinflammation induced by MPTP *in vivo*. CASPASE-1 deficiency significantly reduces motor dysfunctions and dopaminergic neurodegeneration in MPTP-treated mice. These results highlight a mechanism of microglial function that is dependent on CASPASE-1-GSDMD-Pyrpptosis axis and required for PD progression.

## Abbreviations

PD: Parkinson’s disease
MPTP: 1-methyl-4-phenyl-1,2,3,6-tetrahydropyridine
MPP^+^iodide: 1-methyl-4-phenylpyridinium iodide
VX-765: Belnacasan
SN: substantia nigra
GSDMD: Gasdermin-D
NLRP3: NOD-like receptor thermal protein domain associated protein 3
PAMP: Pathogen-Associated Molecular Patterns
DApMP: Damage-Associated Molecular Patterns
TH: Tyrosine hydroxylase
GFAP: Glial Fibrillary Acidic Protein
α-Syn: α-Synuclein
AD: Alzheimer’s disease

## Acknowledgements

We thank the Strong Foundation Project and the Outstanding Young Talents Training Project and the “Da-zhang” scientific research project of Naval Medical Center, the “Qi-hang” scientific research project of Naval Medical University and the Research Project of Shanghai Municipal Health Commission.

## Declarations

## Ethics approval and consent to participate Statement

All animal experiments follow the reach guidelines (https://arriveguidelines.org) Reported. All animal experiments were conducted in accordance with relevant guidelines and regulations and approved by the Laboratory Animal Center of the Naval Medical University, China in conformance with the National Institute of Health Guide for the Care and Use of Laboratory Animals.

## Competing interests

The authors declare that they have no competing interests.

## Authors’ contributions

You Yin, Jicong Du and Tao Huang designed the study. Mengmeng Wang, Yuedong Wang and Tianyi Wang performed the major experiments. Yang Yao and Kaikai Fu were responsible for the collection of data and interpretation in the animal experiments. Mengmeng Wang was responsible for the writing of the manuscript. Mengmeng Wang and You Yin supported fund assistance. All authors read and approved the final manuscript.

## Funding

This study was supported in part by the grants from the Strong Foundation Project [grant number 23M2701] and the Research Project of Shanghai Municipal Health Commission[20224Y0396] and the “Da-zhang” scientific research project [grant number 27M2702] of Naval Medical Center, the “Qi-hang” scientific research project [grant number 27X2701] of Naval Medical University and the Outstanding Young Talents Training Project [grant number 21TPQN2701]. These support the experimental research and data collection of the subject.

## Data Availability Statement

All data were included in the manuscript. Further inquiries can be directed to the corresponding authors.

## Consent for publication

Not applicable.

